# Analysis of mitochondrial genome methylation using Nanopore single-molecule sequencing

**DOI:** 10.1101/2021.02.05.429923

**Authors:** Theresa Lüth, Christine Klein, Susen Schaake, Ronnie Tse, Sandro Pereira, Joshua Lass, Lasse Sinkkonen, Anne Grünewald, Joanne Trinh

**Author notes:** Corresponding author Joanne Trinh, PhD, Institute of Neurogenetics, University of Lübeck, Germany, Ratzeburger Allee 160, BMF Geb. 67, 23562.

## Abstract

The level and the biological significance of mitochondrial DNA (mtDNA) methylation in human cells is a controversial topic. Using long-read third-generation sequencing technology, mtDNA methylation can be detected directly from the sequencing data, which overcomes previously suggested biases, introduced by bisulfite treatment-dependent methods. We investigated mtDNA from whole blood-derived DNA and established a workflow to detect CpG methylation with Nanopolish. In order to obtain native mtDNA, we adjusted a whole-genome sequencing protocol and performed ligation library preparation and Nanopore sequencing. To validate the workflow, 897bp of methylated and unmethylated synthetic DNA samples at different dilution ratios were sequenced and CpG methylation was detected. Interestingly, we observed that reads with higher methylation in the synthetic DNA did not pass Guppy calling, possibly affecting conclusions about DNA methylation in Nanopore sequencing. We detected in all blood-derived samples overall low-level methylation across the mitochondrial genome, with exceptions at certain CpG sites. Our results suggest that Nanopore sequencing is capable of detecting low-level mtDNA methylation. However, further refinement of the bioinformatical pipelines including Guppy failed reads are recommended.

## Introduction

Methylation of the mitochondrial DNA (mtDNA) has been previously studied in various biological contexts^1,2^. In terms of environmental and lifestyle factors, mtDNA methylation changes have been observed to be associated with air pollution and smoking^3–5^. Specific CpG methylation status within different genes of the mtDNA correlates with airborne particulate matter^3,4^. Furthermore, in smokers, methylation levels of particular CpG sites were higher compared to past- and non-smokers^5^. Aside from environmental factors, changes in mtDNA methylation have been reported to be related to neurodegeneration such as Alzheimer’s disease (AD), Parkinson’s disease (PD)^6^ and amyotrophic lateral sclerosis (ALS)^7^. Nuclear genome methylation correlates with age^8,9^, and it has been suggested that mtDNA can behave similarly, as an association between methylation level of two CpG sites within the mtDNA and age has been observed^10^. Given the importance of environmental impact on mtDNA and its association with disease, the investigation of mtDNA CpG methylation is of interest. However, the level of methylation and its biological significance is still under debate. The underlying reason for this debate can be due to methodology.

The most common method to detect mtDNA methylation has been bisulfite treatment. With bisulfite treatment, the characterization of the methylation pattern depends on the resistance of 5-methylcytosine (5-mC) to be converted to uracil. However, the secondary and tertiary structure of intact mtDNA may block cytosines from bisulfite conversion, which influences the outcome of the detected methylation levels^11,12^. Bisulfite conversion-resistant cytosines can lead to an overestimation of mtDNA methylation. To counteract the bias of bisulfite-resistant cytosines, linearization of mtDNA has been recommended^11^. For linearized mtDNA prior to bisulfite treatment, lower levels of methylation have been observed compared to intact mtDNA^12,13^. The low levels of methylation have led to suggestions of this being artifactual meaning an absence of methylation in the mitochondrial genome altogether^12,14^. On the other hand, there has also been supportive evidence for the presence of mtDNA methylation after linearization^15,16^. Furthermore, cytosines in a non-CpG context have been reported to be relatively highly methylated^16,17^.

Using long-read, single-molecule sequencing technologies like Nanopore sequencing^18^, it is possible to overcome limitations of the indirect measurement of methylation with bisulfite treatment, as it can be detected directly from the squiggle signals^19^. Nanopore sequencing enables the direct sequencing of full-length linearized mtDNA reads in contrast to the shorter fragments obtained from second-generation sequencing^20^. To our knowledge, there are currently only two studies that have used Nanopore sequencing to investigate mtDNA methylation^21,22^. Thus, detecting mtDNA methylation with this novel sequencing technology remains to be thoroughly investigated and validated.

Herein, we focus on the exploration of CpG methylation on the mtDNA using Nanopore sequencing. We present a workflow to analyze mtDNA CpG methylation from human blood-derived DNA using a modified library preparation step and analysis pipeline on six individuals. In addition, we compare strand-specific CpG methylation of the mtDNA to the nuclear-encoded *45S* rRNA gene. We then validated our methylation pipeline with a short 897 bp synthetic DNA product, suggesting that highly methylated reads are prone to lower Guppy base-calling Phred quality scores.

## Materials and Methods

### Demographics of individuals

All individuals included in this study provided informed consent and local ethics committee approval at the Research Ethics Board of the University of Luebeck was obtained. In this study, six individuals were included (L-3244, L-13062, L-6581, L-2131, L-2132 and L-2135). Three of the included individuals are patients with PD who also have *Parkin* mutations, with two being biallelic mutation carriers (L-3244 and L-13062) and one heterozygous carrier (L-6581) (Supplementary Table 1). The other three are healthy control subjects (L-2131, L-2132 and L-2135). Movement disorder specialists performed the clinical assessments of patients. The genetic screening for *Parkin* was performed with Sanger Sequencing and MLPA from blood-derived DNA, as previously described^23^. DNA extraction from blood was performed with the QIAGEN blood DNA midi kit following the manufacturer’s instructions.

#### Library preparation and Nanopore sequencing

##### Whole genomes

DNA concentration of the blood-derived DNA was quantified with Qubit fluorometric quantification using the dsDNA BR Assay kit (Thermo Fisher Scientific). Library preparation of 1.5 μg input DNA from the six DNA samples was performed with the ligation sequencing kit (SQK-LSK109) following a Nanopore whole-genome sequencing protocol (premium WGA). To conserve DNA methylation, whole-genome amplification with random primers was omitted from this established protocol (the intended first step of the premium WGA protocol).

##### Synthetic DNA

To further validate the workflow, two types of commercially available 897 bp synthetic DNAs (Zymo Research, D4505) were used where all cytosines were modified to 5-mC or unmodified. From these two synthetic DNAs, we created three different dilutions: 1) only methylated DNA (100% methylation), 2) 1:1 dilution of methylated and unmethylated DNA (50% methylation), and 3) only unmethylated DNA (0% methylation).

DNA concentration was quantified as described above and 1 μg input DNA was used. The library was prepared using the Nanopore ligation sequencing kit (SQK-LSK109) and multiplexed with the native barcoding expansion (EXP-NBD104).

##### Sequencing

Nanopore sequencing was performed with the MinION using the R9.4.1 flow cells. The flow cell was primed and loaded according to the manufacturer’s instructions. For base-calling, MinKNOW (version 20.10.6) on the MinIT with the integrated Guppy (version 4.2.3) was used. The Guppy base-calling software is available for Nanopore community members (https://community.nanoporetech.com). We aimed to obtain 10 Gigabases (Gb) of data before stopping the run which lasted ~24 hours.

### Data analysis

#### DNA methylation analysis

Base-called Nanopore reads were aligned to the reference sequence (i.e., mitochondrial genome, hg38 assembly), using Minimap2 (version 2.17). DNA methylation in a CpG dinucleotide context was called with Nanopolish (version 0.13.2), a software tool that uses a *Hidden Markov Model* (HMM)^24^. Nanopolish reports the log-likelihood ratio for each observed event (i.e., CpG site within a *k-mer* sequence). The methylation frequency (MF) was calculated with the default threshold of the log-likelihood ratio. Thus, with a log-likelihood ratio >2 the CpG site was classified as methylated and with a value <-2 the site was classified as unmethylated. In addition, the split-group parameter was used, which provides the MF of each detected CpG site in the reference sequence separately. The MF describes the proportion of reads that support methylation at the given CpG site. To investigate the methylation of the synthetic DNA, alignment, methylation calling, and calculation of MF were performed. The D5405 reference sequence provided by the manufacturer was used. Reads that passed Guppy calling, as well as failed reads, were analyzed.

#### Plus- and minus-strand methylation comparison

To explore heavy-strand (H-strand) and light-strand (L-strand) methylation, methylation calls were stratified by plus- and minus-strand and then the MF was calculated. The plus-strand represents the L-strand and the minus-strand the H-strand.

In order to compare mtDNA to a nuclear-encoded gene, we used the *45S rRNA* gene (13 kb core region of the 45S cluster 5). There are multiple copies of the *45S rRNA* gene in the human genome, comparable in size and coverage to the mtDNA. DNA methylation of the *45S rRNA* gene was stratified by plus- and minus-strand to allow comparison to the H- and L-strand of mtDNA.

#### Coverage and mtDNA methylation analysis

In order to investigate the relationship between coverage and MF, different subsets of the obtained sequencing data (i.e., Fast5 and FastQ files) with increasing coverage were used to detect methylation. The mean coverage and MF of each data set were calculated.

#### Statistical analysis

For statistical analysis, GraphPad Prism software (version 9) was used. To compare median MF between plus- and minus-strand, the non-parametric Mann-Whitney U test for pairwise comparison was performed. For correlation analyses, the non-parametric Spearman correlation was used. All p-values were exploratory and not corrected for multiple testing.

## Results and Discussion

We investigated mtDNA methylation in six blood-derived samples from three patients with PD carrying Parkin mutations and three control subjects. We obtained a mean of 10 Gb of data and a read length of 5.4kb with whole-genome Nanopore sequencing. Alignment to the mitochondrial genome (hg38) resulted in a mean coverage of 269X (SD=±58.59).

### Mitochondrial DNA methylation

Overall, we detected low-level methylation (mean MF±SD=0.060±0.054) of mtDNA across the six samples. However, there were 32 unique CpG sites, where an MF higher than 0.2 was detected (range of MF=0.2-0.793, Figure 1). These sites were located in the following genes: *12S rRNA* (chrM:707, 807, 1022, 1261, 1413, 1560), *16S rRNA* (chrM:1748, 1785, 2475, 2583, 2842, 2914, 2941, 3008), *ND1* (chrM:3405), *ND2* (chrM:4711, 4918), *COI* (chrM: 6052, 6241, 6688), *COII* (chrM: 8253), *ATPase6/8* (chrM:8646), *ND4* (chrM:11590, 12123) *tRNA-Histidine* (chrM:12190), *ND5* (chrM:12455), *tRNA-Glutamic acid* (chrM:14696), tRNA-Threonine (chrM:15929) and in the non-coding D-loop (chrM: 314, 16127, 16359, 16410). Each individual had at least five sites that showed >0.2 MF. *ND1* (chrM:3405) consistently showed >0.2 MF in all samples (range of MF=0.333-0.403).

**Figure 1.**
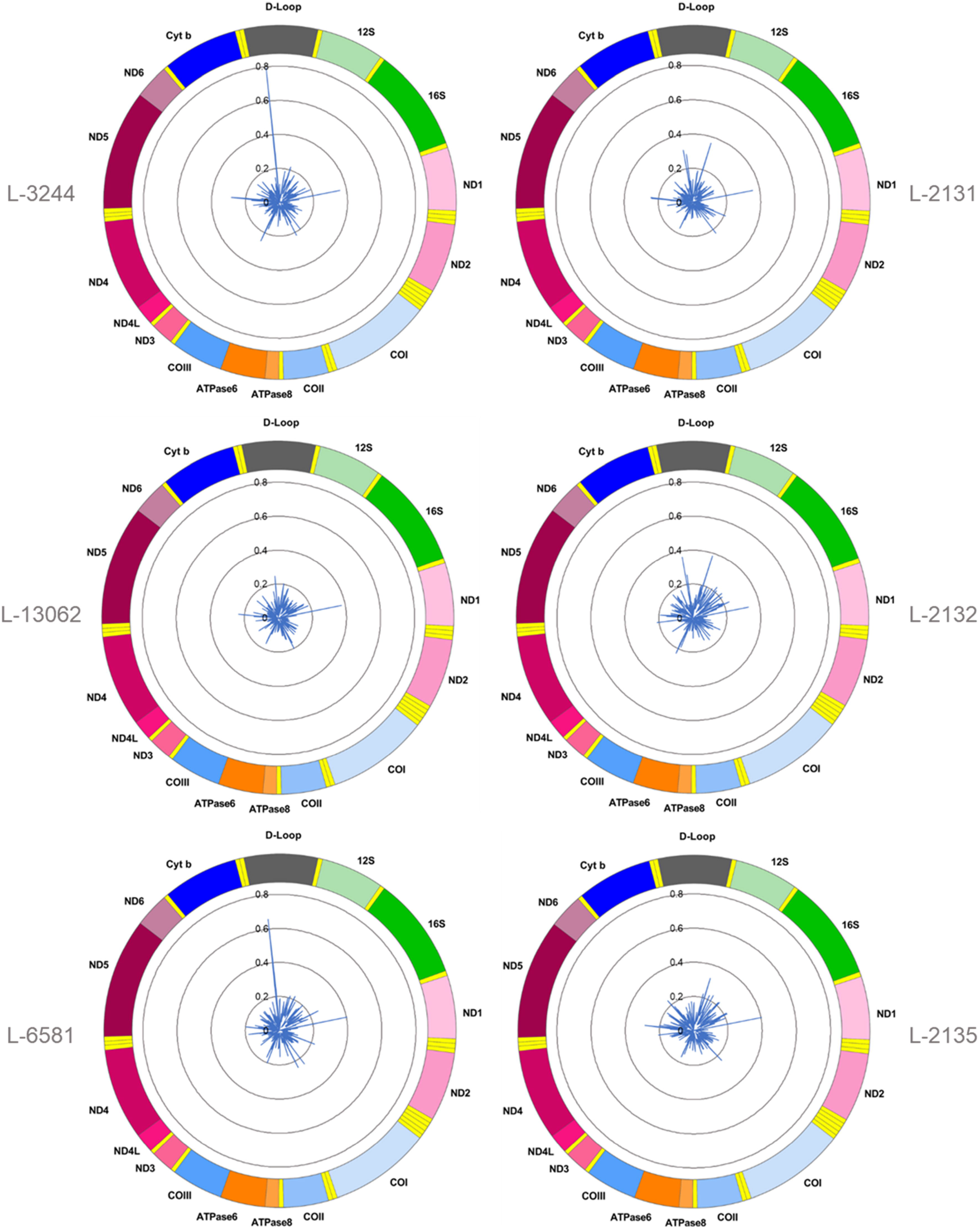
Mitochondrial CpG methylation frequency detected by Nanopore sequencing in six individuals. Radar chart showing mitochondrial DNA methylation from six blood-derived DNA samples detected by Nanopolish. Methylation frequency is indicated by blue spikes. 12S=Small subunit rRNA; 16S=Large subunit rRNA; ND1=NADH dehydrogenase, subunit 1; ND2=NADH dehydrogenase, subunit 2; COI=Cytochrome c oxidase, subunit 1; COII=Cytochrome c oxidase, subunit 2; ATPase8=ATP synthase, subunit 8; ATPase6=ATP synthase, subunit 6; COIII=Cytochrome c oxidase, subunit 3; ND3=NADH dehydrogenase, subunit 3; ND4L=NADH dehydrogenase, subunit 4L; ND4=NADH dehydrogenase, subunit 4; ND5=NADH dehydrogenase, subunit 5; ND6=NADH dehydrogenase, subunit 6; Cyt b=Cytochrome b; D-Loop=Displacement loop

### Heavy- and light-strand methylation differences

We next explored differences in methylation of the H- and L-strand. The H-strand has a higher purine content (G and A) compared to the L-strand and encodes 12 out of the 13 mitochondrial-encoded proteins^25^. As the reference sequence of the mitochondrial genome represents the L-strand, Nanopolish reports the L-strand as the plus-strand and the H-strand as the minus-strand^24^.

The MF detected from the mtDNA plus- and minus-strands was positively correlated (Spearman’s r=0.3683, Spearman’s exploratory p-value<0.0001, Figure 2A). For comparison, the *45S rRNA* gene was used as a nuclear-encoded comparison, since it has comparable size and coverage to the mtDNA (size=13 kb, mean coverage=623X). The overall correlation between plus-strand and minus-strand methylation was stronger in the *45S* rRNA gene (Spearman’s r=0.7482, Spearman’s exploratory p-value<0.0001, Figure 2B) compared to the mtDNA.

**Figure 2.**
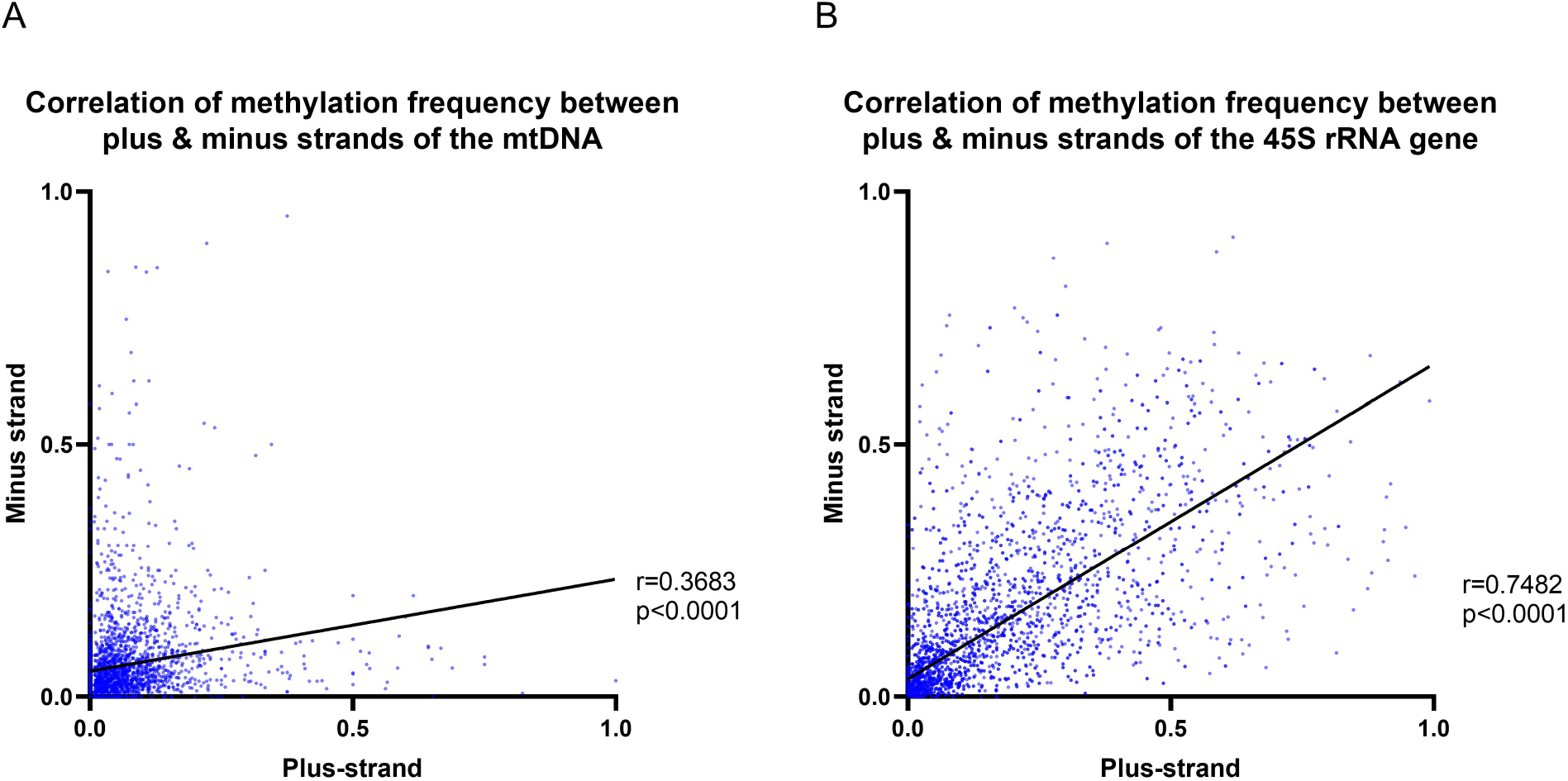
Mitochondrial and 45S rRNA CpG methylation comparison by strand. A) Correlation between the methylation frequency of the mitochondrial plus- and minus-strand (plus-strand=light-strand, minus-strand=heavy-strand). Data of the six blood-derived samples combined. B) Correlation between the methylation frequency detected from the 45S rRNA gene plus- and minus-strand. Data of the six blood-derived samples combined. r=Spearman’s rank correlation coefficient, p=Spearman’s exploratory p-value

In the mtDNA, certain CpG sites consistently showed high methylation on one strand. For example, the CpG site within the *ND1* gene (at position chrM:3,405), showed consistently higher MF on the minus-strand (mean MF=0.801) compared to the plus-strand (mean MF=0.084) in all six individuals. Similarly, a CpG site within the *45S rRNA* gene (at position 45S: 10654) showed consistently higher MF on the minus-strand (mean MF=0.645) compared to the plus-strand (mean MF=0.047).

Although there was a trend for differences in the median MF between mtDNA plus- and minus-strands in five individuals (Supplementary Figure 1A, Mann Whitney U-test p<0.0270), the difference was not consistently in the same direction and effect sizes were small (difference in median MF<0.020). Furthermore, the p-values were not corrected for multiple testing and the Mann-Whitney U test for pairwise comparison was only performed to explore a potential trend towards higher or lower methylation levels in one of the strands. Similar to the mtDNA, the CpG sites in *45S rRNA* showed a trend for differences in the median MF between plus- and minus-strands in five individuals (Supplementary Figure 1B, Mann Whitney U-test p<0.0101, difference in median MF<0.063). Methodologically, there was no preferential sequencing for either of the strands (Supplementary Figure 1C/D, Mann Whitney U-test p>0.05). Given the low effect of differences between L- and H-strands, we cannot suggest strand-specific CpG methylation in mtDNA.

### Relationship between the coverage, number of called sites and methylation frequency

The mean MF for the six individuals was at ~0.06 with >100X coverage (Figure 3A). The relationship between coverage and MF of all detected individual CpG sites did not correlate (Spearman’s r=0.0009, Spearman’s exploratory p-value>0.05, Figure 3B). However, the number of called sites and MF was negatively correlated (Spearman’s r=-0.3114, Spearman’s exploratory p-value<0.0001, Figure 3C). The number of called sites represents how often a given CpG site was evaluated as methylated or unmethylated by Nanopolish^24^. Likewise, a negative association between coverage and the detected methylation frequency was shown with whole-genome bisulfite sequencing^12^ and there were speculations that bisulfite conversion-resistant cytosines were the culprit. However, the detection of mtDNA methylation from the Nanopore sequencing data does not depend on bisulfite conversion and we observed a plateau of the mean MF after 100X coverage. Furthermore, with all generated sequencing data included, we did not observe a negative association between the coverage and the MF of detected CpG sites. On the other hand, we observed a negative correlation between the number of called sites and the detected MF, which could be due to the general low-level methylation of the mtDNA.

**Figure 3.**
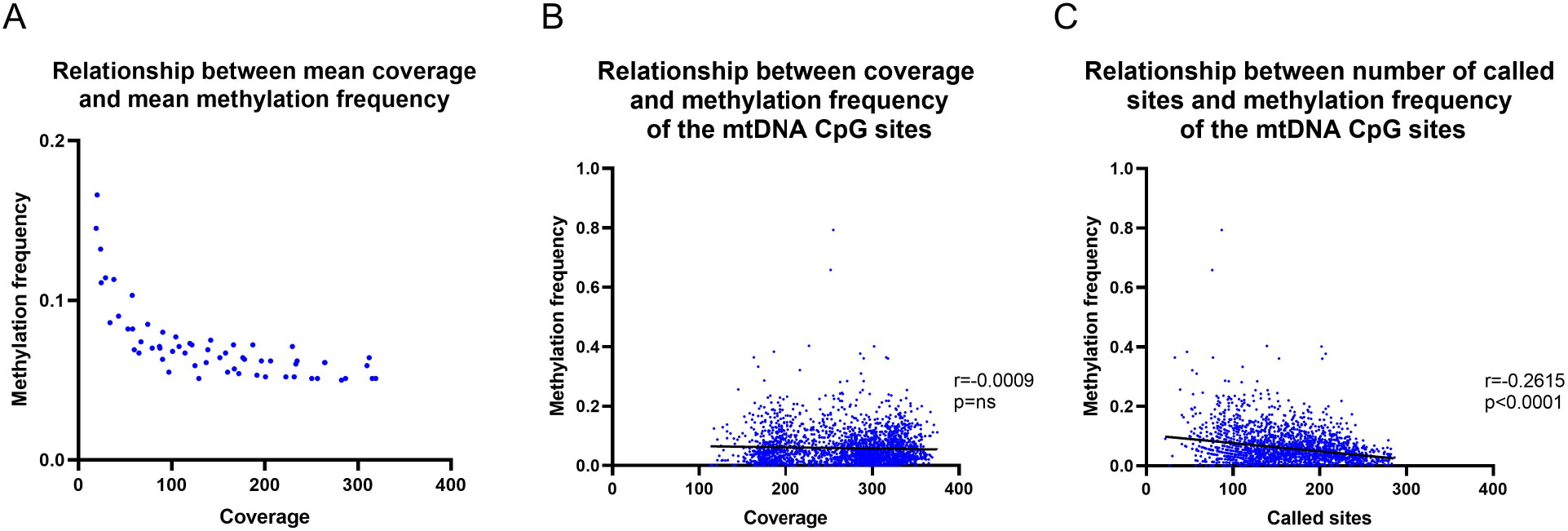
Relationship between coverage and methylation frequency. A) Overall relationship between the mean coverage and mean methylation frequency of different subsections of the obtained sequencing data. B) Overall relationship between the coverage of each individual CpG and the methylation frequency. C) Overall relationship between the number of called sites of each CpG and the methylation frequency. r=Spearman’s rank correlation coefficient, p=Spearman’s exploratory p-value

### Synthetic DNA

To further validate the workflow, we performed Nanopore sequencing of an 897 bp synthetic methylated and unmethylated control DNA sample at different dilution ratios (0%, 50% and 100%). We obtained a mean of 14Mb data and a read length of 963 bp. As the three samples were multiplexed for sequencing, barcodes with a length of 40 bp were added on both sides of the DNA fragments, which led to the higher mean read length of 963 bp compared to the 897 bp reference sequence. Alignment resulted in a mean coverage of 5018X (SD=±424X).

### Synthetic DNA methylation

The CpG methylation of three samples with a ratio of 0%, 50% or 100% of methylated synthetic DNA molecules was investigated (Figure 4A). In the 0% methylated sample, an MF=0.021 (SD±0.042) was detected similar to the 50% methylated sample (mean MF±SD=0.022±0.038). In the 100% methylated sample, the detected methylation was higher (mean MF±SD=0.262±0.141). Particularly at CpG sites (e.g. D4505:195-350), the detected methylation level in the 0% methylated sample was higher than zero (range of MF=0.014-0.193), which could indicate false-positive calls. On the other hand, the detected methylation levels were overall lower than expected in the 50% and 100% methylated samples and the detected methylation levels in the 0% and 50% methylated samples were almost indistinguishable.

**Figure 4.**
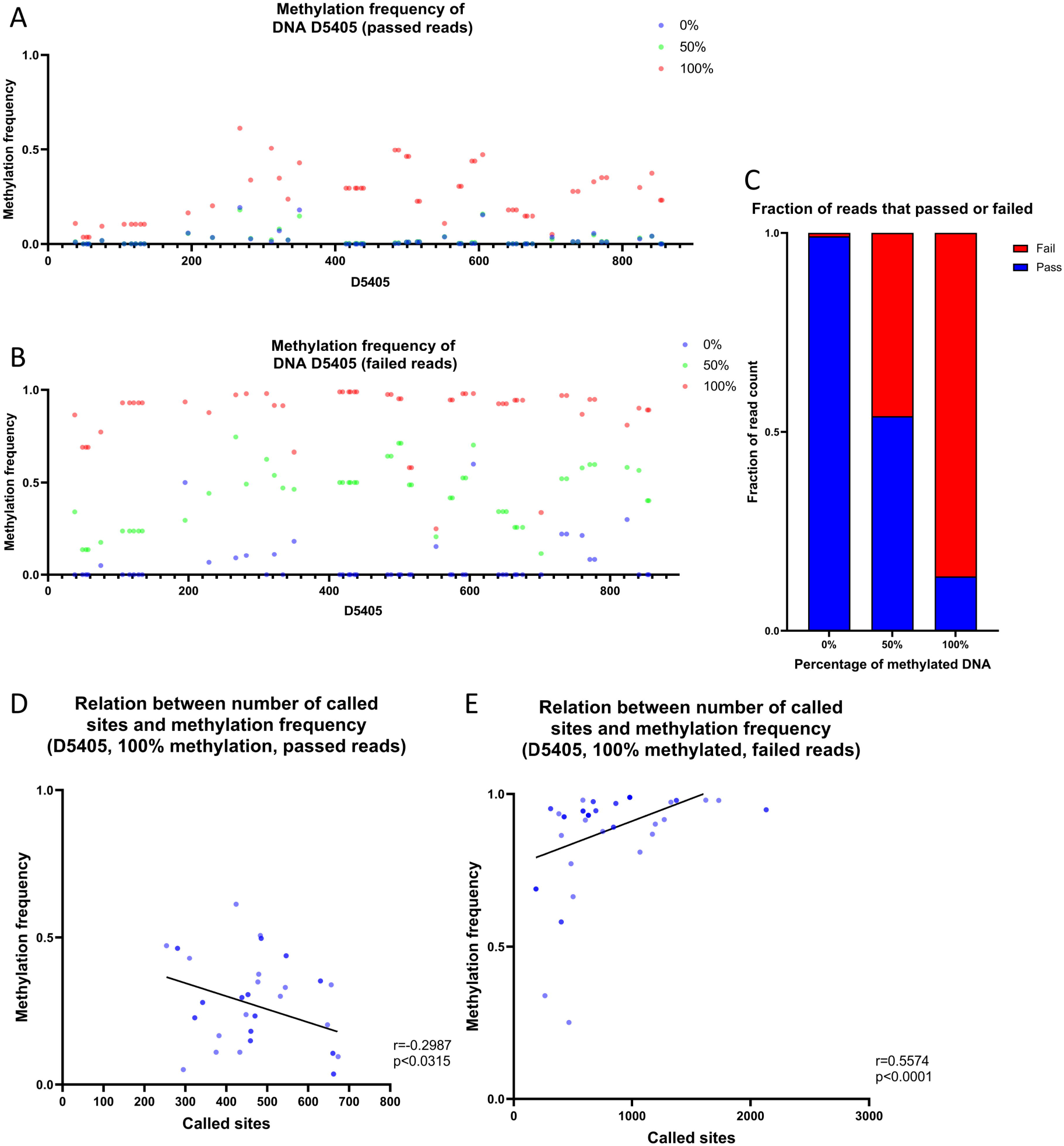
Analysis of synthetic DNA for the validation of the data analysis pipeline. A) Methylation frequency detected by Nanopolish from three samples of synthetic DNA with different proportions of methylated DNA (0%, 50% and 100%), only reads that passed Guppy quality-threshold were included in the analysis. B) Methylation frequency detected by Nanopolish from three samples of synthetic DNA with different proportions of methylated DNA (0%, 50% and 100%), only reads that failed Guppy quality-threshold were included in the analysis. C) Stacked bar plot of the fraction of reads that passed (blue) or failed (red) the Guppy quality-threshold, stratified by the proportion of methylated reads in the sample. D) Relationship between the number of called sites and methylation frequency in the sample with 100% methylated reads, only reads that passed Guppy quality-threshold were included in the analysis. E) Relationship between the number of called sites and methylation frequency in the sample with 100% methylated reads, only reads that failed Guppy quality-threshold were included in the analysis. r=Spearman’s rank correlation coefficient, p=Spearman’s exploratory p-value

### Investigation of Guppy failed reads

To investigate the possibility of preferential sequencing of unmethylated reads and methylated reads which do not pass the quality threshold during base-calling, we analyzed Guppy failed reads. During the Guppy base-calling, the software reports Phred quality scores (qscores) of each read, which indicates the accuracy of the base-calling. Reads with low qscores are classified as failed. Exploring these failed reads only, we detected low methylation in the 0% methylated sample (mean MF±SD=0.057±0.123) and higher levels in the 50% methylated sample (mean MF±SD=0.430±0.168) and the 100% methylated sample (mean MF±SD=0.880±0.156). Thus, the detected methylation was overall higher in Guppy failed reads compared to Guppy passed reads and showed a better representation of the dilution ratios (Figure 4B). As the detected MF in the 0% methylated sample was higher in the failed reads compared to the passed, there is the possibility of even more false-positive calls.

Including both the passed and failed reads, the proportion of failed reads was n=55/6550 in the 0% methylated sample, n=26068/56577 in the 50% methylated sample and n=44505/51555 in the 100% methylated sample (Figure 4C), which further shows that methylated reads are prone to lower Guppy qscores. Furthermore, there was a negative correlation between the number of called sites and the detected MF from passed reads in the 100% methylated sample (Spearman’s r=-0.2987, Spearman’s exploratory p-value=0.0315, Figure 4D) compared to a positive correlation when the MF was detected from the failed reads (Spearman’s r=0.5574, Spearman’s exploratory p-value<0.0001, Figure 4E). In the 50% methylated sample, there was a similar change in the direction of the relationship between the number of called sites and MF in the passed reads (Spearman’s r=-0.6078, Spearman’s exploratory p-value<0.0001) compared to the failed reads (Spearman’s r=0.3048, Spearman’s exploratory p-value=0.0280). In contrast, the 0% methylated sample showed a negative correlation between the number of called sites and the MF in the passed reads (Spearman’s r=-0.6452, Spearman’s exploratory p-value=<0.0001), but there was no correlation in the failed reads (Spearman’s r=0.2310, Spearman’s exploratory p-value=ns).

Although the methylation patterns from the failed reads were more representative of the corresponding dilution ratios of methylated DNA, a higher probability for sequencing errors will be present in the failed reads.

### Reanalysis of mitochondrial methylation including Guppy failed reads

To consider the results obtained from the investigation of the synthetic DNA samples, mtDNA methylation was reanalyzed, including Guppy passed and failed reads. We obtained a mean coverage of 280X (SD=±62X) of the mitochondrial genome, which was on average 11X more coverage compared to just including the passed reads. Since the number of failed reads were low, the overall change in the MF was marginal (Supplementary Table 2). However, the methylation “peak” in the D-loop, for samples L-3244 (MF=0.793) and L-6581 (MF=0.688) was even higher after the inclusion of the failed reads (L-3244: MF=0.797, L-6581: MF=0.695). Thus, reanalysis of highly methylated CpG sites within mtDNA may be of importance.

### Review of literature on mtDNA Nanopore sequencing

We performed a literature search for the terms “mtDNA” and “Nanopore” and found 36 publications, seven of which performed Nanopore sequencing of human mtDNA^21,22,26–30^ (Supplementary Figure 3). In these studies, the coverage of the mitochondrial genome ranged from 60.4X^21^ to over 10,000X^22^ (Supplementary Table 3). In the latter case, extraction of mtDNA from a subcellular fraction was performed before Nanopore sequencing, which enhanced the fraction of sequenced reads that mapped to the mitochondrial genome. Two out of the six publications investigated mtDNA methylation in the context of cancer research^21,22^. Although in our study, we included patients with PD and *Parkin* mutations, we did not further investigate the significance of mtDNA methylation differences in the context of PD. Possible differences in mtDNA methylation between patients and control subjects in this small sample set are speculative and future studies to thoroughly investigate mtDNA methylation in PD are warranted. For the published studies in cancer cells, Nanopolish was used to call methylation from the Nanopore sequencing data^21,22^. In addition to Nanopolish, one study used Guppy in combination with Medaka^22^. Low-level mtDNA methylation in these two studies has been described^21,22^, albeit one study evaluated only three specific sites. We also detected overall low-level mtDNA methylation in blood-derived DNA, in concordance with reported literature^21,22^.

### Strengths and limitations

In our study, we present an optimized protocol for the investigation of mtDNA CpG methylation. We obtained high coverage of the mtDNA as well as the nuclear-encoded *45S rRNA* gene and the synthetic DNA samples. The mean coverage of the mtDNA ranged from 177X to 320X across individuals. Linearization of mtDNA has been recommended to counteract the influence of secondary and tertiary structure in bisulfite-dependent methods^11–13^. Thus, we performed T7 Endonuclease I digestion of whole-cell DNA for single-molecule sequencing. Still, the sample size of our study was small with six included individuals and our analysis was limited to DNA methylation in a CpG context. As previous studies have shown that there is non-CpG methylation present in the mtDNA^16,31^, future workflows should be expanded to include non-CpG methylation, as well. We were also unable to measure the exact amount of linearized mtDNA contained in the sample, which can have an impact on the reflection of MF. Given these limitations, we investigated two different types of genetic material in addition to the 16.5 kb mitochondrial genome to validate our workflow i.e, an 897bp synthetic DNA sample and the native 13kb *45S rRNA* nuclear-encoded gene. Lastly, all experiments and analyses were performed in the same institute to reduce batch effects.

### Conclusion

In conclusion, our results suggest the applicability of Nanopore sequencing for the investigation of mtDNA methylation. Overall, we detected low-level CpG methylation of mtDNA with exceptions for certain sites.

Our results suggest that highly methylated DNA molecules were more likely to fail Guppy base-calling and therefore bioinformatical pipelines that take Guppy failed reads into account are recommended.

## Acknowledgments

We acknowledge Dr. Aurelien Ginolhac for his review of the manuscript and insightful comments. Funding has been obtained from the German Research Foundation (“ProtectMove”; FOR 2488, GR 3731/5-1; SE 2608/2-1; KO 2250/7-1), the Luxembourg National Research Fund within the ATTRACT (“Model-IPD”, FNR9631103) and INTER programs (“ProtectMove”, INTER/DFG/19/14429377), the Personalized Medicine Consortium (PMC) of Luxembourg (PUMP-PRIME grant), the European Community (SysMedPD), the Canadian Institutes of Health Research (CIHR), Alexander von Humboldt Foundation, and the Joachim Herz Stiftung.

