## Supplementary Figure 1 for "Analysis of mitochondrial genome methylation using Nanopore single-molecule sequencing"

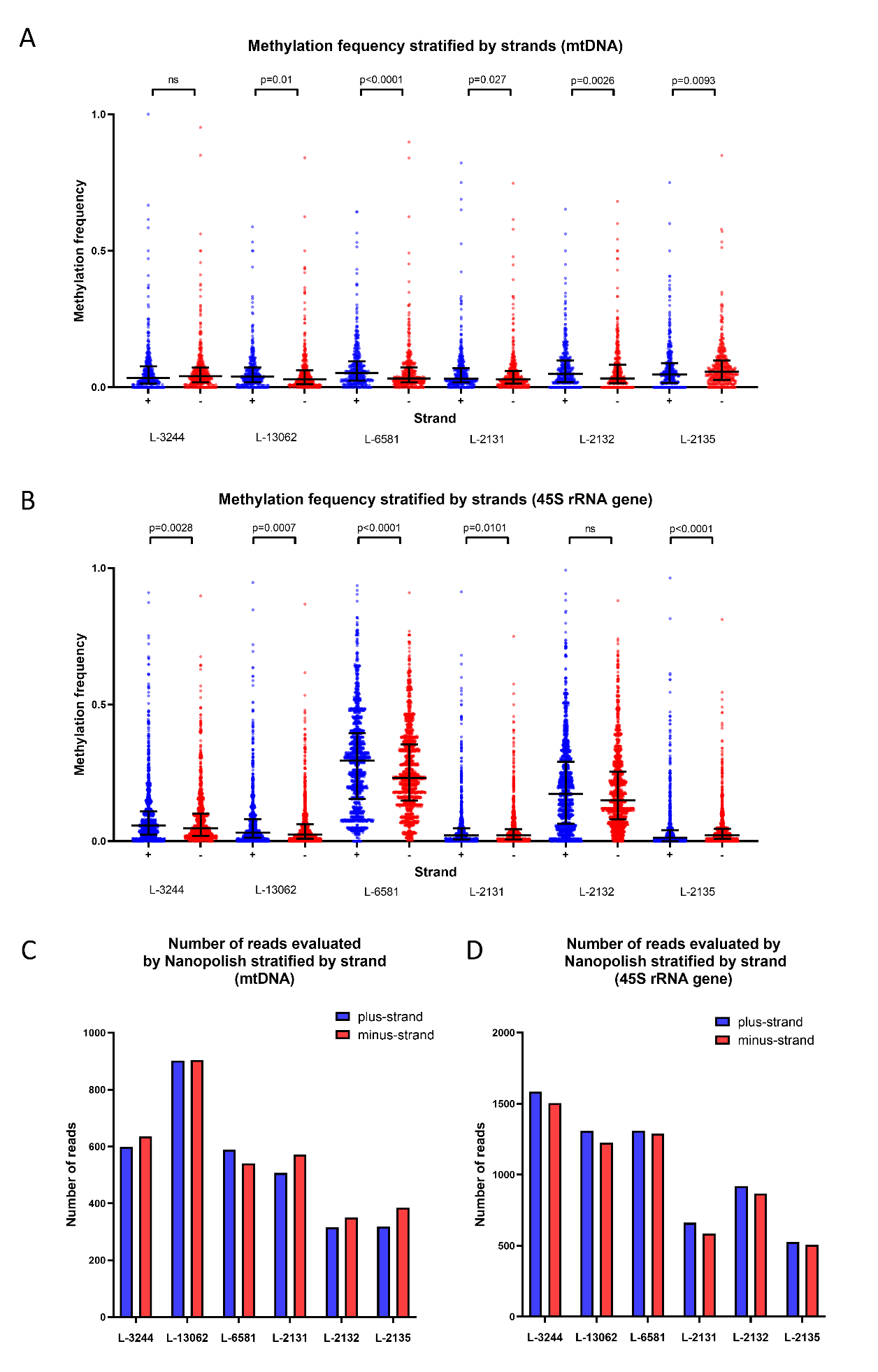

**Supplementary Figure 1. Exploration of DNA CpG methylation stratified by strand.** A/B) Scatter plot of the methylation frequency stratified by strands from the mtDNA or the 45S rRNA gene . Bars indicating median and inter quartile range (IQR), p-value=Mann Whitney U-test performed for pairwise comparisons. C/D) Bar plot showing number of reads mapped to the mitochondrial genome or the 45S rRNA gene and evaluated by Nanopolish, stratified by strands.
