## Supplementary Figure 2 for "Analysis of mitochondrial genome methylation using Nanopore single-molecule sequencing"

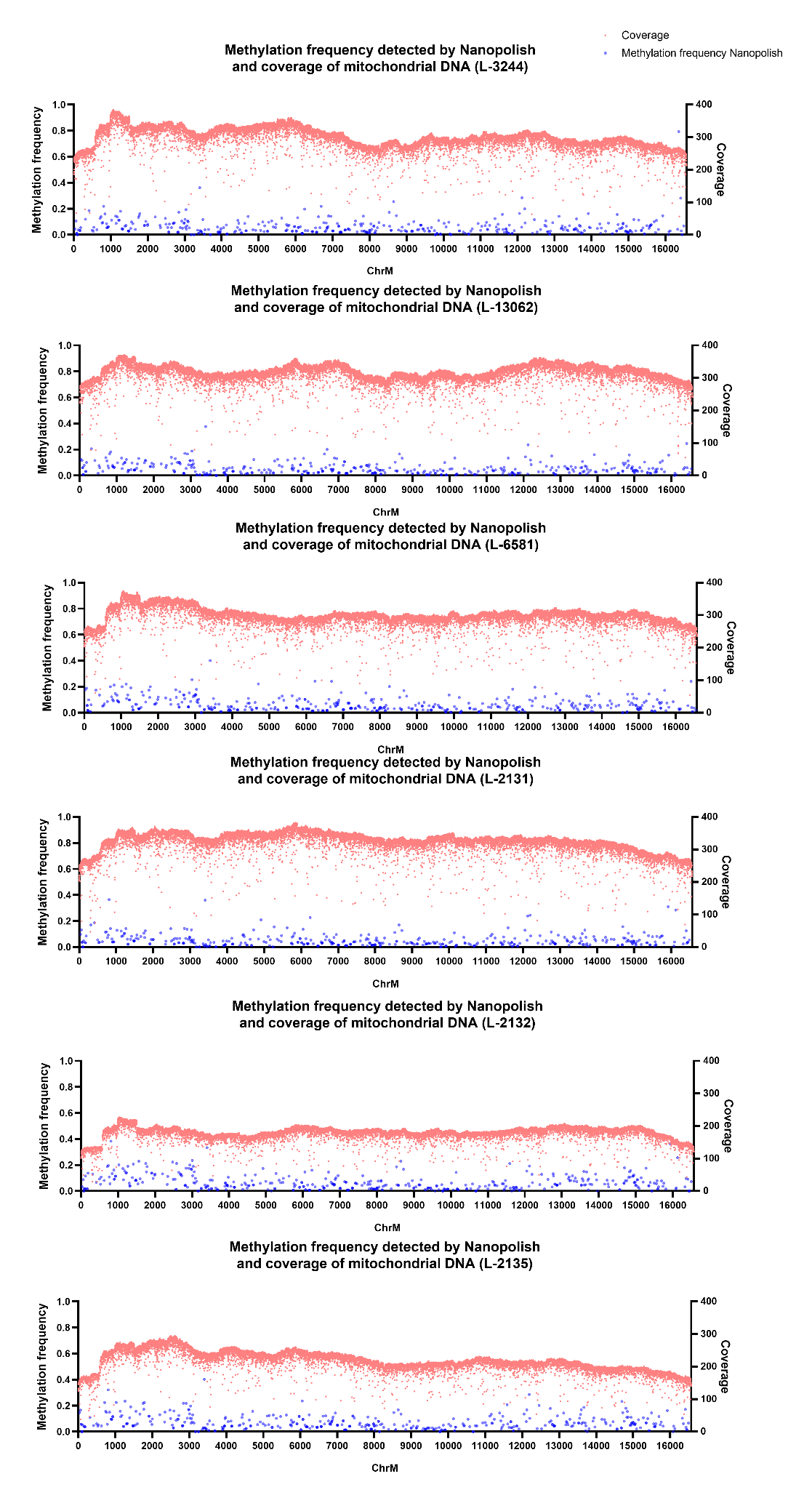

**Supplementary Figure 2. Mitochondrial CpG methylation frequency and coverage detected by Nanopore sequencing in six individuals**. The methylation frequency of the mitochondrial DNA from six blood-derived DNA samples, detected by Nanopolish. Methylation frequency is indicated by blue dots and coverage by red dots and the x-axis indicates the positions in the mitochondrial genome (hg38).
