## Supplementary Figure 3 for "Analysis of mitochondrial genome methylation using Nanopore single-molecule sequencing"

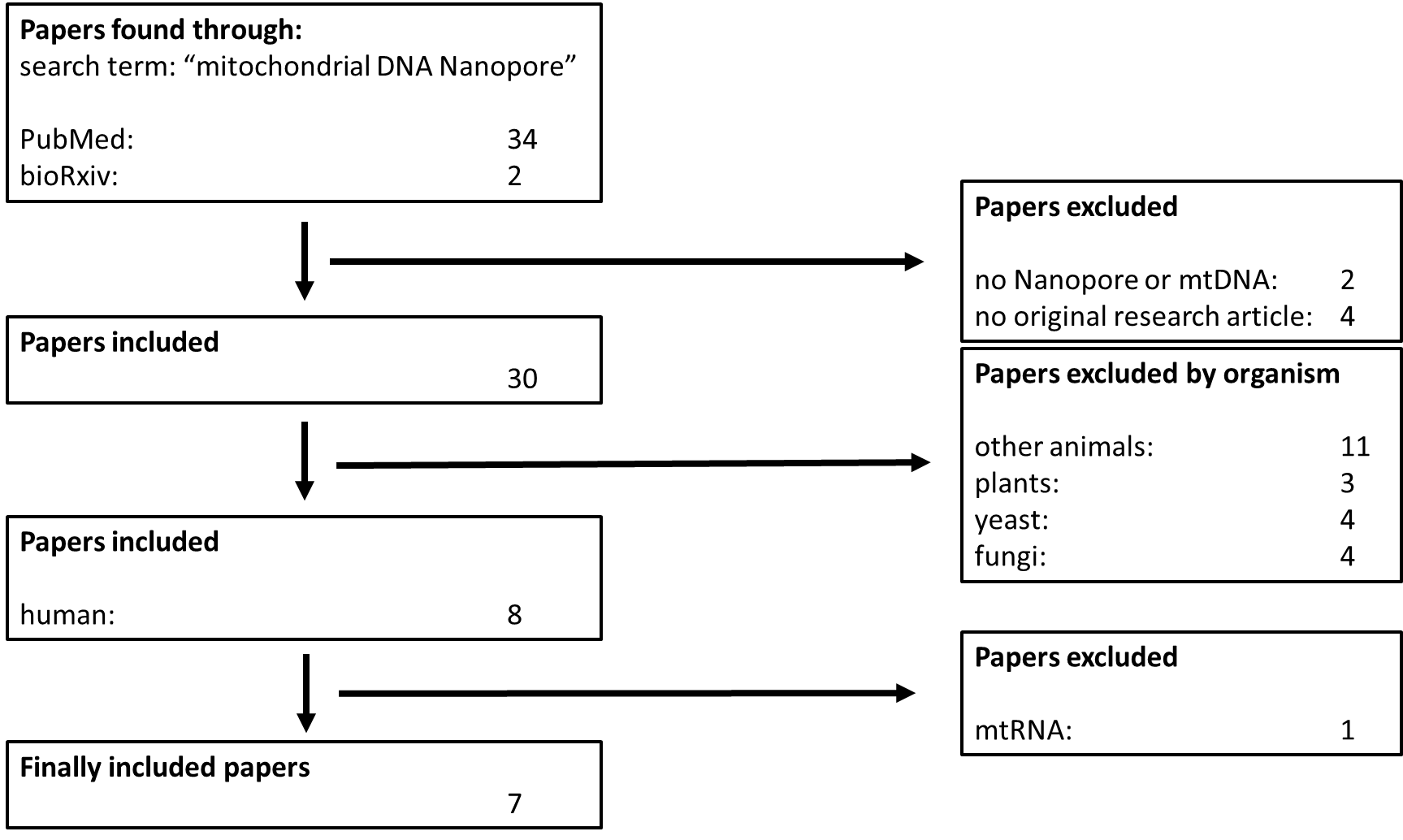

**Supplementary Figure 3. Flowchart summarizing literature search for articles on mitochondrial DNA and Nanopore sequencing.** The Search term was: “mitochondrial DNA Nanopore” and the date of the literature search was: January 29th, 2021.
