## Supplementary Table 1 for "Analysis of mitochondrial genome methylation using Nanopore single-molecule sequencing"

|  | L-3244 | L-13062 | L-6581 | L-2131 | L-2132 | L-2135 |
| --- | --- | --- | --- | --- | --- | --- |
| AAE | 49 | 17 | 69 | 57 | 35 | 39 |
| AAO | 36 | 11 | 58 | n.a. | n.a. | n.a. |
| Gender | Female | Female | Female | Male | Female | Male |
| Parkin variant I  - Protein level identifier  - cDNA level identifier  - Genomic location hg(38) on chr6 | - p.Arg275Trp  - c.823C>T  - 6:161785820 | - p.Met1?  - c.2T>C  - 6:162727667 | - p.Pro437Leu  - c.1310C>T  - 6:161350187 | n.a. | n.a. | n.a. |
| Parkin variant I  - Deletion  - cDNA level identifier | - exon 1 del  - c.(?_-103-1)_(7+1_8-1)del | - exon 11 del  - c.(1167+1_1168-1)_(1285+1_1286-1)del | n.a. | n.a. | n.a. | n.a. |

**Supplementary Table 1.** Demographics of study participants

n.a.=Not applicable, AAE=Age at examination, AAO=Age at onset, Parkin variant I/II=Description of Parkin variants of the compound heterozygous (L-3244, L-13062) or heterozygote (L-6581) mutation carriers with PD, del=Deletion
