## Supplementary Table 2 for "Analysis of mitochondrial genome methylation using Nanopore single-molecule sequencing"

**Supplementary Table 2**. Methylation frequency of certain CpGs in the mitochondrial DNA before and after including reads that failed Guppy quality-threshold.

| Gene (position) | Guppy quality filtering | L-3244  MF (called sites) | L-13062  MF (called sites) | L-6581  MF (called sites) | L-2131  MF (called sites) | L-2132  MF (called sites) | L-2135  MF (called sites) | Average MF (average called sites) |
| --- | --- | --- | --- | --- | --- | --- | --- | --- |
| 12S  (807) | Pass | 0.218 (87) | 0.167 (90) | 0.193 (83) | 0.364 (77) | 0.383 (47) | 0.321 (53) | 0.274 (73) |
|  | Pass/Fail | 0.213 (89) | 0.176 (91) | 0.198 (86) | 0.359 (78) | 0.383 (47) | 0.315 (54) | 0.274 (74) |
| ND1  (3405) | Pass | 0.361 (169) | 0.377 (207) | 0.400 (195) | 0.360 (203) | 0.333 (111) | 0.403 (139) | 0.372 (171) |
|  | Pass/Fail | 0.349 (175) | 0.364 (209) | 0.392 (199) | 0.357 (207) | 0.330 (112) | 0.394 (142) | 0.364 (174) |
| tRNA-Thr  (15925) | Pass | 0.145 (62) | 0.157 (83) | 0.091 (77) | 0.310 (58) | 0.364 (33) | 0.163 (43) | 0.205 (59) |
|  | Pass/Fail | 0.143 (63) | 0.155 (84) | 0.101 (79) | 0.300 (60) | 0.364 (33) | 0.163 (43) | 0.204 (60) |
| D-Loop  (16359) | Pass | 0.793 (58) | 0.062 (96) | 0.688 (80) | 0.072 (97) | 0.097 (62) | 0.062 (64) | 0.296 (76) |
|  | Pass/Fail | 0.797 (59) | 0.077 (104) | 0.695 (82) | 0.071 (99) | 0.094 (64) | 0.062 (65) | 0.299 (79) |
| D-Loop  (16410) | Pass | 0.281 (57) | 0.246 (65) | 0.254 (59) | 0.125 (64) | 0.091 (22) | 0.171 (41) | 0.195 (51) |
|  | Pass/Fail | 0.293 (58) | 0.293 (70) | 0.254 (59) | 0.123 (65) | 0.080 (25) | 0.171 (41) | 0.202 (53) |

Gene (position)=Gene name and mitochondrial DNA start position (0-based), MF=Methylation frequency, which is the proportion of reads that support the methylated state at that CpG site, Called sites=Number of reads that could be classified as methylated or unmethylated by Nanopolish at that CpG site, 12S=Small subunit rRNA, ND1=NADH dehydrogenase, subunit 1 (complex I), tRNA-Thr=Transfer RNA for Threonine, D-Loop=Displacement Loop
