## Supplementary Table 3 for "Analysis of mitochondrial genome methylation using Nanopore single-molecule sequencing"

| **Author**  **Supplementary Table 3**. Overview of literature search for articles on mitochondrial DNA and Nanopore sequencing. | **Year** | **Coverage of mtDNA** | **Methylation** | **Tools for methylation calling** | **Sequencing strategy (Kit, Flow cell number, machine)** | **Sample**  **Type (sample size)** | **Disease investigated** | **Enrichment methods** | **Conclusions** | **Reference** |
| --- | --- | --- | --- | --- | --- | --- | --- | --- | --- | --- |
| Aminuddin | 2020 | 60.4X | yes | Nanopolish | native mtDNA,  PCR amplicons  (SQK-LSK108, FLO-MIN107, MinION) | Cell line (SAS, H103) | Oral squamous cell carcinoma | Two PCR amplicons for the entire mitochondrial genome (8kb) | - Lower cisplatin sensitivity could be caused by genetic and epigenetic changes of mitochondrial genome | ^1^ |
| Goldsmith | 2020 | >10,000X | yes | Nanopolish  Guppy + Medaka | native mtDNA  (SGK-RBK004, FLO-MIN160, MinION) | Cell line (HepaRG, HEK293T),  liver tissue | Cancer | Subcellular fractionation, targeted nanopore sequencing | - Low-level of strand specific CpG methylation, higher methylation in tissue compared to cell lines | ^2^ |
| Georgieva | 2020 | 7,000X | no | - | CRISPR/Cas9  (SQK-LSK109, FLO-MIN106D, MinION) | Embryonic stem cells (C9012) | - | High Pure PCR template preparation kit | - ONT can detect 11 different thymidine analogs and determine replication rates | ^3^ |
| Zascavage | 2019 | 295-813X | no | - | native mtDNA, PCR amplicon  (SQK-RAD001, MKI vR9, MinION) | Blood, cell line (HL-60) | - | PCR amplicon of entire mitochondrial genome | - Direct mtDNA sequencing from native DNA is a reliable alternative to approaches using PCR-enriched libraries | ^4^ |
| Carter | 2017 | 68X | no | - | native DNA  (NSK007, FLO-MIN106, MinION) | Cell line (HAP1) | - | CsgG-based sequencing | - ONT CsgG-based sequencing may be useful for complex genomes | ^5^ |
| Bi | 2020 | NA | no | - | Amplified mtDNA (SQK-LSK109, FLO-MIN106, MinION) | Mouse oocytes, Human | NARP/Leigh syndrome | PCR amplicon for the entire mitochondrial genome | - iMiGseq of full mtDNA is good for ultra-senstive variant detection, complete haplotyping and unbiased evaluation of heteroplasmy level | ^6^ |
| Lindberg | 2016 | ~200X | no | - | PCR amplicon (-, FLO-MAP003, MinION) | Blood | - | 2 PCR amplicons for the entire mitochondrial genome | - Hybrid assembly of MiSeq and MinION can accurately recontruct full mitochondrial genomes | ^7^ |
